# Excluding loci with substitution saturation improves inferences from phylogenomic data

**DOI:** 10.1101/2021.08.28.457888

**Authors:** David A. Duchêne, Niklas Mather, Cara Van Der Wal, Simon Y.W. Ho

## Abstract

The historical signal in nucleotide sequences becomes eroded over time by substitutions occurring repeatedly at the same sites. This phenomenon, known as substitution saturation, is recognized as one of the primary obstacles to deep-time phylogenetic inference using genome-scale data sets. We present a new test of substitution saturation and demonstrate its performance in simulated and empirical data. For some of the 36 empirical phylogenomic data sets that we examined, we detect substitution saturation in around 50% of loci. We found that saturation tends to be flagged as problematic in loci with highly discordant phylogenetic signals across sites. Within each data set, the loci with smaller numbers of informative sites are more likely to be flagged as containing problematic levels of saturation. The entropy saturation test proposed here is sensitive to high evolutionary rates relative to the evolutionary timeframe, while also being sensitive to several factors known to mislead phylogenetic inference, including short internal branches relative to external branches, short nucleotide sequences, and tree imbalance. Our study demonstrates that excluding loci with substitution saturation can be an effective means of mitigating the negative impact of multiple substitutions on phylogenetic inferences.

---

One of the key steps in phylogenomics is to identify a suitable set of loci for reconstructing the evolutionary history of a group of organisms. The inference of phylogenetic trees from nucleotide sequences can be misled by a number of factors. For example, the sequences might contain too little information if they have evolved too slowly (Yang 1998; Klopfstein et al. 2017). On the other hand, high evolutionary rates will increase the probability of multiple substitutions occurring at the same nucleotide sites, leading to a phenomenon known as substitution saturation (Brown et al. 1982; Mindell and Honeycutt 1990; Philippe and Forterre 1999; Philippe et al. 2011). Even when the best-fitting model of nucleotide substitution is used, saturation can cause the phylogenetic method to produce inaccurate estimates of the tree topology and branch lengths. Therefore, an important step in experimental design for phylogenomic analysis is to identify the loci with excessive evolutionary rates that might mislead phylogenetic inference (Philippe et al. 2011).

One popular approach for exploring phylogenetic informativeness is to identify the loci that have experienced substitution saturation (Philippe and Forterre 1999). A widely used approach for this purpose is to compare the sites in a sequence alignment with those expected under conditions of complete saturation. Following information theory, the test takes the pattern of nucleotide frequencies at a fully saturated site to follow a multinomial distribution with maximum entropy (Xia et al. 2003). The critical values for this test are the entropy values above which the estimates of the tree topology and branch lengths are likely to be inaccurate.

In an extensive simulation study, Xia et al. (2003) described the behaviour of the entropy test across a range of phylogenetic conditions of overall evolutionary rate, amounts of data, and tree imbalance. However, there is a wide range of factors that can interact to mislead phylogenetic inference, such as substitution model underparameterization (Revell et al. 2005; Sullivan and Joyce 2005) or the presence of long terminal branches (Klopfstein et al. 2017; Dornburg et al. 2019). Some of these factors are rapidly gaining recognition in phylogenomics research, in which identifying sources of bias can be crucial for obtaining a reliable estimate of the phylogeny (Reddy et al. 2017; Mai and Mirarab 2018). Therefore, understanding the sensitivity of tests of saturation to a broad range of phenomena in empirical data can improve practice in phylogenomics.

Data sets in phylogenomics are typically composed of many alignments of non-recombining regions of the genome (loci). The phylogenetic information signal in each locus is often summarized using its ‘gene tree’. Tests of substitution saturation can be used to select loci for phylogenetic analysis (e.g., Han and Ro 2005; Dávalos and Perkins 2008; Liu et al. 2014). This form of data filtering ultimately aims to maximize the signal of the true phylogenetic relationships in the data, also known as the historical signal. There has been growing interest in data-filtering methods for phylogenomics (Molloy and Warnow 2018; Richards et al. 2018; Bravo et al. 2019). However, the effectiveness of these methods, such as using tests of saturation, remains to be explored in depth.

In this study, we describe the performance of a common test of saturation, and evaluate the impact of saturation in a broad range of empirical phylogenomic data sets. We also describe a novel approach to examining substitution saturation that focuses exclusively on phylogenetically informative sites. This approach greatly ameliorates the negative influence of slowly evolving sites on the measurement of overall base composition, which is central to the performance of the entropy-based tests of saturation. Using two simulation studies, we first aim to identify the characteristics of sequence alignments to which the tests are sensitive, including a high rate of substitution. In our second simulation study, we explore a broad continuum of evolutionary scenarios to examine the power of the tests to identify loci with amounts of substitution saturation that are likely to mislead estimates of phylogeny and branch lengths. We then evaluate the degree of substitution saturation in 36 phylogenomic data sets, and investigate the link between saturated loci and amount of discordance in phylogenetic signal across sites. Our results suggest that saturation can affect large portions of phylogenomic data sets, and that testing for saturation is an effective approach to identify loci with poor historical signal in phylogenomic studies.

## Materials and Methods

### Test of expected entropy

The degree of substitution saturation in a nucleotide sequence alignment can be described using a measure of entropy (Xia et al. 2003). The entropy of a distribution is defined as the average information content in a given sample. The information content measures the level of surprise expected when we encounter a particular outcome. Systems with high entropy are more unpredictable and disorderly. In the context of phylogenetics, entropy is highest at full saturation, when noise has overridden the historical signal. In neutrally evolving sequences under full saturation, every site might be expected to have nucleotide frequencies equal to those of the whole alignment. The nucleotide that occurs in each sequence at a site is assumed to be independent of the nucleotides in other sequences, so that the vector of nucleotides at each site can be modelled as a single draw from a multinomial distribution, where the underlying probabilities are equal to the nucleotide frequencies. The entropy of a multinomial sample with *n* observations, *k* categories, and a vector *p* of probabilities for each of the categories is:

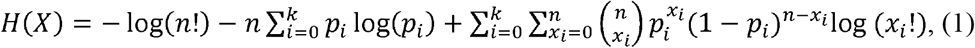

Therefore, the expected entropy of a sequence alignment under full saturation can be calculated using the number of taxa (*n*) and the overall nucleotide frequencies (*p*). Meanwhile, the information content at an observed site is based on the counts of nucleotides at that site:

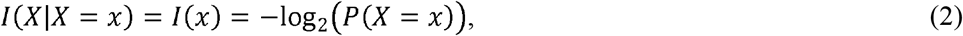

Where *P*(*X* = *x*) is the density of a sample from the multinomial distribution, given that the probabilities are the overall nucleotide frequencies in the sequence alignment. Since the entropy is the expected value of the information content, we can calculate the entropy of our sample by taking the average of the information at each site. A test of saturation can then be performed by taking the estimates of entropy across alignment sites and comparing this with the sample entropy expected under full saturation.

Substitution saturation can have negative impacts on phylogenetic inference even before the historical signal is completely eroded (Ho and Jermiin 2004), so a classic test of the significance of the distance (*t*) between the distribution of observed site information against the value of maximal entropy will provide only limited information about phylogenetic inferences. Instead, a critical value for the *t*-statistic test can be derived from simulations (Xia et al. 2003). The critical value *t_crit_* can then be used for testing the hypothesis that the empirical value of *t_obs_* is far from maximal entropy, a situation in which we would expect a negligible impact of substitution saturation on phylogenetic inference. Our formulation of this test is slightly different from that of Xia et al. (2003), who tested whether *t_obs_* is significantly smaller than *t_crit_*. We propose testing whether *t_obs_* is significantly smaller than full entropy, using *t_crit_* as the critical value of the test. Our test considers the variance in the information content across sites more explicitly, but the two tests can be expected to produce similar results.

Existing implementations of the test of saturation take the expected entropy to depend on the base frequencies of the sequence alignment. However, this test can have problematic behaviour in several scenarios, such as when sequences are subject to selective constrains or when sites vary in evolutionary rate. Such variation in rates is common in empirical data sets: among codon positions in protein-coding sequences or across sites in ultraconserved elements (UCEs) and their flanking regions. Therefore, we assessed the performance of the test based exclusively on phylogenetically informative sites (i.e., parsimony-informative sites with at least two nucleotides, where at least two of these occur in at least two sequences). This avoids the worst effects of constant sites on the test in data matrices with small numbers of taxa, small numbers of sites, or both. The focus of the test on phylogenetically informative sites will have a diminishing effect on the test as the size of the data matrix increases, and is expected to have a null effect under infinite sites.

The guidelines require revision so that the test can be implemented more effectively for empirical data. Very small sequence alignments can pose challenges for the reliability of the test, due to the disproportionate effect of small numbers of outlier sites. Similarly, alignments with large numbers of slow-evolving sites and an overrepresentation of a particular nucleotide type will be highly sensitive to single outlier sites. Therefore, a test of saturation using the entropy *t*-statistic is unlikely to perform well on small sequence alignments or in which nucleotide frequencies are highly uneven.

### The entropy t-statistic and phylogenetic inference

We used a simulation study to examine the circumstances under which substitution saturation is a significant problem that can be addressed in empirical data, relative to other causes of misleading inferences. We simulated the evolution of nucleotide sequences along trees with variable numbers of taxa (8, 32, 128, and 512) to generate data sets with three different sequence lengths (250, 500, and 1500 nucleotides). Starting with trees that were fully symmetric and in which all branches had equal length, we varied several conditions that can affect the performance of phylogenetic methods, including: the number of substitutions along each branch of the tree (0.05, 0.25, 0.45, and 0.65); tree imbalance (fully balanced tree versus fully imbalanced tree); ratio of the sum of internal to the sum of external branch lengths, or stemminess (0.1, 0.5, and 0.9; Fiala and Sokal 1985); and the substitution model that generated the data (the matched Jukes-Cantor model, JC; or the more complex GTR+Γ model). Simulations under GTR+Γ focused on examining the impact of model underparameterization, and were made with transition parameters drawn from a Dirichlet distribution (all values of *α* = 5) and gamma-distributed rates across sites with *α* = 1. In addition, we simulated sequence evolution under the conditions above but including a proportion of constant sites (0, 0.25, 0.5, 0.75), as is common in several empirical data types. Our simulations produced 100 sequence alignments under each combination of scenarios and under either the JC or the GTR+Γ substitution model. We performed phylogenetic analyses of each data set using the JC substitution model and maximum-likelihood optimization in IQ-TREE (Nguyen et al. 2015).

Phylogenetic accuracy in each scenario was calculated as the unweighted and normalized Robinson-Foulds distance (Robinson and Foulds 1981; Penny and Hendy 1985) between the inferred tree and the ‘true’ tree used for simulation in each analysis. We also made a comparable calculation with branch lengths, by taking the difference between the estimated and true tree length, and dividing this difference by the true tree length. The outcome is the proportion error in estimated tree length. In addition, we recorded the entropy *t*-statistics as calculated on all alignment sites and on informative sites only.

We tested the hypothesis that each factor that we varied in our simulations had an impact on phylogenetic inference and on the two entropy *t*-statistics. We tested four linear regression models in which the response variables were, in turn, the topological distance between the estimated and true trees, the difference in tree length between the estimated and true trees, and each of the two *t_obs_* values of the saturation test (calculated on all sites or only on informative sites). The explanatory variables were the six fixed factors that we varied in our simulations (number of taxa, alignment length, tree length, degree of tree imbalance, stemminess, and substitution model used for simulation). In addition, we considered the total number of variable sites in the alignment, and all of the two-way interactions between the main effects. The resulting *p*-values were corrected for multiple comparisons using false discovery rates.

### Diagnostic ability of the entropy t-statistic

In a second simulation study, we characterized the performance of the two entropy *t*-statistics for identifying cases in which substitution saturation is misleading phylogenetic inference. We explored a similar parameter space as in our first simulation study, but treated several of the factors as continuous. In each simulation, we sampled values from uniform distributions for mean branch length U(0.01, 0.65), stemminess U(0.1, 0.9), and sequence length U(250, 2000). In each simulation, we also randomly sampled the number of taxa (8, 32, 128, and 512), whether the tree was imbalanced or balanced, a portion of the sites to remain constant U(0, 0.8), and the model of nucleotide substitution used for sequence evolution (JC or GTR+Γ). We repeated 10^5^ times the sampling of parameters, simulation of sequence evolution, phylogenetic inference in IQ-TREE under a JC model, and the calculation of the two *t*-statistics of entropy (all sites versus informative sites only). This analysis allowed us to explore the performance of the test statistics along a gradient of scenarios and to identify appropriate critical values for interpreting the best-performing test statistic.

We quantified the diagnostic ability of the two entropy *t*-statistics across the simulation conditions by using receiver operating characteristic (ROC) curves to show the relationship between rates of true positives and false positives. Across a traverse of the values of the test statistic calculated in simulations, we considered positives to be cases in which the topology of the inferred phylogeny was different from that of the true tree. For each ROC curve, we chose the critical values as those that maximized the difference between the numbers of true positives and false positives, such that they provide the greatest power for discriminating positives from negatives. The code used to perform simulations is freely available online (github.com/duchene/entropy_saturation_test).

We also explored the performance of the entropy *t*-statistics in identifying cases in which substitution saturation is likely to have misled estimates of branch lengths. Branch lengths are continuous variables, so an arbitrary critical value has to be used to determine the target positives for the test. We determined a positive as being cases in which estimated tree length was at least 50% greater or smaller than the true tree length.

Based on the results of our simulations, we propose critical values for assessing saturation using the entropy test statistic calculated on phylogenetically informative sites. Past research has indicated that test statistics often cannot be interpreted using a single, universally applied critical value, because the power of any given test can vary with sample size and other features of the data (e.g., Xia et al. 2003; Duchêne et al. 2017). Following previous work, we propose critical values that depend on the number of taxa and on sequence length. Specifically, we performed a multiple regression of the critical value chosen under each simulation condition on the square root of the corresponding number of taxa and sequence length as explanatory variables. Regression models can then be used to predict critical values across any number of taxa and sequence length. The saturation test is implemented in the free software PhyloMAd (Duchêne et al. 2018b; github.com/duchene/phylomad) and reports the test results, a diagnosis based on the new critical values, and estimates of expected false-positive and true-positive rates.

### Signal of substitution saturation in phylogenomic data

The prevalence and impact of substitution saturation was examined in a range of phylogenomic data sets from the literature (Table 1). We obtained 36 data sets that varied widely in taxonomic range, size, and sequence type. The data sets spanned multiple orders of magnitude in their number of taxa and number of genomic regions included. They also included a range of data types that we broadly describe as exon, intron, ultraconserved element, or anchored-enriched region data. While these data types have some overlap, they are different in the data targeted by researchers (e.g., coding gene regions versus the informative regions flanking UCEs), which potentially allows useful distinctions to be made.

**Table 1.** Phylogenomic data sets tested for substitution saturation in this study. Data sets were examined as done in previous studies, such that each locus was analysed independently.

| Taxon | Number of loci<sup>a</sup> | Number of taxa per locus | Data type/genomic region<sup>b</sup> | Mean run time of saturation test per locus (s) | Source reference |
| --- | --- | --- | --- | --- | --- |
| Stinging wasps (Aculeata) | 807 | 140–183 | UCE | 0.273 | Branstetter et al. 2017 |
| Bilateria metazoans (Bilateria) | 424 | 50–75 | Exon | 0.076 | Cannon et al. 2016 |
| Laurasiatherian mammals (Laurasiatheria) | 10258 | 8–23 | Intron | 0.096 | Chen et al. 2017 (a) |
| Laurasiatherian mammals (Laurasiatheria) | 3637 | 5–23 | Intron | 0.118 | Chen et al. 2017 (b) |
| Amniote vertebrates (Amniota) | 1145 | 10 | UCE | 0.035 | Crawford et al. 2012 |
| Marsupial mammals (Marsupialia) | 1494 | 38–45 | Exon | 0.098 | Duchêne et al. 2018a |
| Butterflies (Papilionoidea) | 350 | 144–205 | Exon | 1.086 | Espeland et al. 2018 |
| Ray-finned fishes (Actinopterygii) | 489 | 5–27 | UCE | 0.040 | Faircloth et al. 2013 |
| North American tarantulas (*Aphonopelma*) | 581 | 63–83 | Anchored | 0.179 | Hamilton et al. 2016 (a) |
| Spiders (Araneae) | 326 | 22–34 | Anchored | 0.107 | Hamilton et al. 2016 (b) |
| North American mygalomorph spiders (Euctenizidae) | 403 | 18–25 | Anchored | 0.085 | Hamilton et al. 2016 (c) |
| Ray-finned fishes (Actinopterygii) | 1101 | 105–298 | Exon | 1.095 | Hughes et al. 2018 |
| Cichlid fishes (Cichlidae) | 533 | 57–149 | Anchored | 0.927 | Irisarri et al. 2018 |
| Birds (Aves) | 8293 | 42–52 | Exon | 0.154 | Jarvis et al. 2014 (a) |
| Birds (Aves) | 8287 | 42–52 | Exon | 0.117 | Jarvis et al. 2014 (b) |
| Birds (Aves) | 2515 | 39–52 | Intron | 0.540 | Jarvis et al. 2014 (c) |
| Gobioid fishes (Actinopterygii: Gobioidae) | 570 | 43 | Exon | 0.133 | Kuang et al. 2018 |
| Iguanas (Phrynosomatidae) | 580 | 4–11 | UCE | 0.052 | Leaché et al. 2015 |
| Flowering plants (Angiosperms) | 370 | 29–35 | Anchored | 0.089 | Léveillé-Bourret et al. 2018 |
| Mosses (Bryophyta) | 105 | 68–146 | Exon | 0.290 | Liu et al. 2019 |
| Birds (Neoaves) | 1539 | 17–33 | UCE | 0.041 | McCormack et al. 2013 |
| Songbirds (Passeri) | 515 | 106 | UCE | 0.191 | Moyle et al. 2016 |
| Acorn ants (*Temnothorax*) | 2091 | 44–50 | UCE | 0.124 | Prebus 2017 |
| Birds (Aves) | 259 | 164–200 | Anchored | 0.750 | Prum et al. 2015 |
| Snakes (*Storeria*) | 322 | 70–90 | Anchored | 0.389 | Pyron et al. 2016 |
| Seed plants | 1308 | 38 | Exon | 0.089 | Ran et al. 2018 (a) |
| (Gymnosperms) |  |  |  |  |  |
| Seed plants<br>(Gymnosperms) | 1308 | 38 | Exon | 0.082 | Ran et al. 2018 (b) |
| Seed plants<br>(Gymnosperms) | 1308 | 38 | Exon | 0.093 | Ran et al. 2018 (c) |
| Harvestmen spiders<br>(Ischirospalidoidea) | 672 | 5 | Exon | 0.037 | Richart et al. 2016 (a) |
| Harvestmen spiders<br>(Ischirospalidoidea) | 653 | 5 | Exon | 0.034 | Richart et al. 2016 (b) |
| Harvestmen spiders<br>(Ischirospalidoidea) | 672 | 5 | Exon | 0.034 | Richart et al. 2016 (c) |
| Ferns<br>(Monilophyta) | 2385 | 52–73 | Exon | 0.118 | Shen et al. 2018 |
| Squamate reptiles<br>(Squamata) | 4175 | 18–34 | UCE | 0.054 | Streicher and Wiens 2017 |
| Squamate reptiles<br>(Squamata) | 44 | 98–167 | Exon | 0.810 | Wiens et al. 2012 |
| Decapod<br>crustaceans<br>(Decapoda) | 105 | 57–94 | Exon | 0.849 | Wolfe et al. 2019 |
| Squamate reptiles<br>(Squamata) | 52 | 98–2378 | Anchored | 1.086 | Zheng and Wiens 2016 |

<sup>a</sup> For some studies, only a subset of the data were examined because of numerical problems (possibly caused by missing data) or file-format difficulties (such as those caused by unusual characters).
<sup>b</sup> “Exon” and “Intron” data sets refers to gene regions, either protein-coding or intronic, respectively; “UCE” refers to ultraconserved elements (Faircloth et al. 2012); “Anchored” refers to anchored-enriched regions (Lemmon et al. 2012).

Our proposed test of substitution saturation was performed separately for each locus in each data set, where ‘locus’ is defined as in the studies from which we obtained the data. Our analyses focused on the test on informative sites only, given that a test on complete alignments has inferior performance under most conditions (see *Results*). We only explore loci with a minimum of 30 phylogenetically informative sites, and in which none of the nucleotide frequencies is greater than 0.5. These bounds ameliorate the severe impact that outlier sites can have on the estimates of the *t*-statistic. For each data set, we compared loci with high versus low risk of saturation in terms of their total numbers of sites, numbers of informative sites, GC content, and phylogenetic inferences. Phylogenetic analyses were performed for each locus using maximum likelihood in IQ-TREE (Nguyen et al. 2015) with the best-fitting substitution model from the GTR+Γ family. As a metric of branch support, we calculated the approximate likelihood-ratio test (aLRT), which assesses the agreement across sites regarding the maximum-likelihood resolution (Guindon et al. 2010). From the inferences for each locus we extracted the mean branch support across branches, mean estimated branch lengths, and stemminess (Fiala and Sokal 1985).

## Results

### The entropy t-statistic and phylogenetic inference

Our simulations show that substitution saturation, simulated as long tree lengths, is one of the primary factors misleading phylogenetic inference (Fig. 1). However, several other factors are also important obstacles to phylogenetic inference, including tree imbalance, large numbers of taxa, low relative lengths of internal to external branches (stemminess), and high complexity in the true substitution process compared with the model used for inference (model underparameterization; Fig. 1).

**Figure 1.**
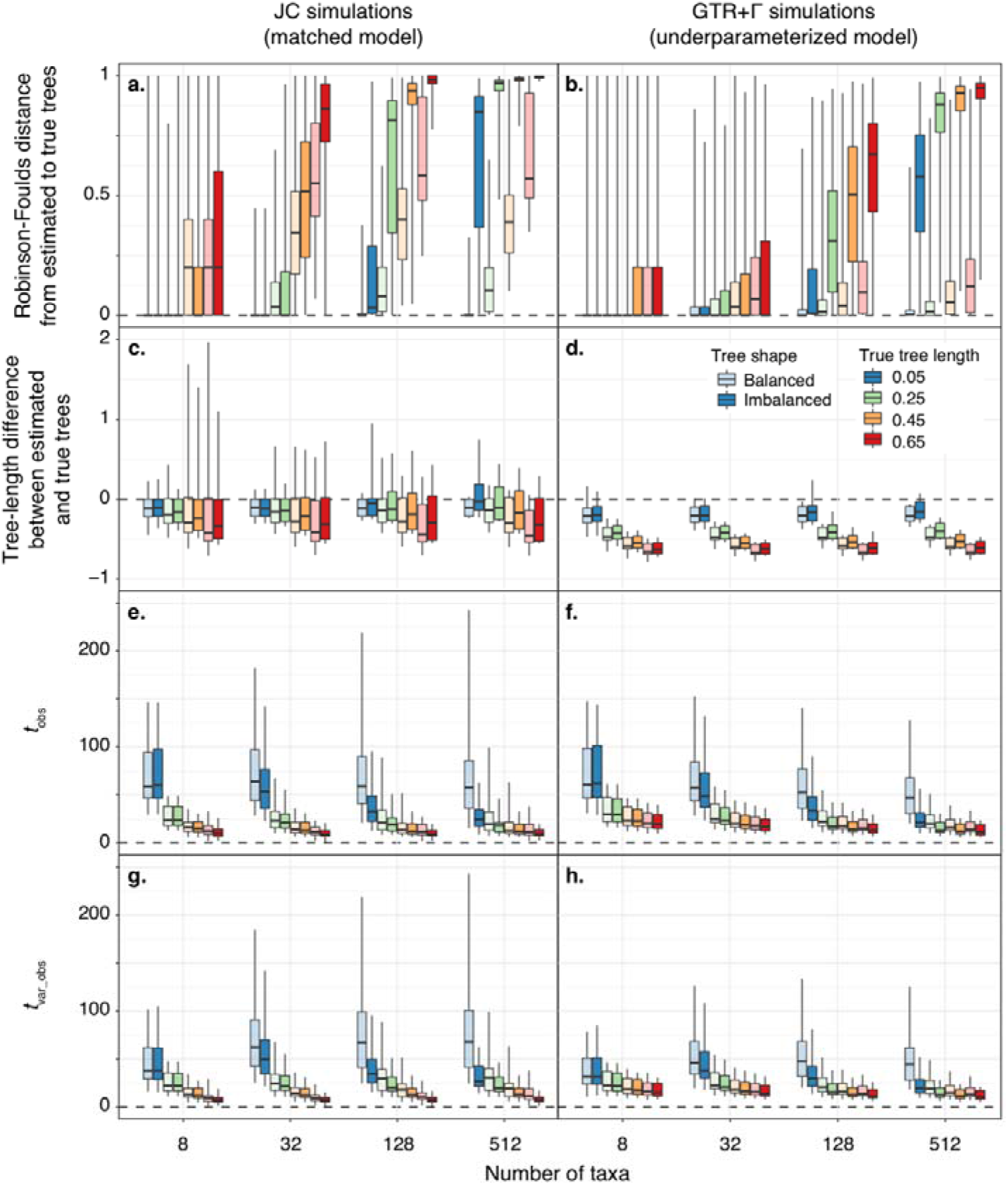
The effect of the most influential factors in simulations on the performance of phylogenetic inference and the entropy *t*-statistic. The JC model of nucleotide substitution was used for all analyses, such that the model is matched in simulations under the JC model (a, c, e, and g), and underparameterized in simulations under the GTR+Γ model (b, d, f, and h). The performance of phylogenetic inference was measured as (a, b) the unweighted Robinson-Foulds distance between the estimated and true trees, and (c, d) the difference in tree length between estimated and true trees, calculated as ((estimated tree length – true tree length) / true tree length). Performance is compared with the entropy *t*-statistic as calculated on all alignment sites (c, d; *t_obs_*) and on phylogenetically informative sites only (c, d; *t_var_obs_*). Boxplot whiskers cover the full range of values in each scenario.

Our regression models highlight the widespread interactions between factors in misleading phylogenetic inferences. For example, substitution saturation is highly misleading to topology inference when tree imbalance is high and trees have large numbers of taxa (Fig. 1a, b). Accuracy in the inferred tree topology was best explained by the two-way interactions between number of taxa and phylogenetic imbalance of the true tree (*t*-value = 218.971; Supplementary Table S1), that between phylogenetic imbalance and stemminess of the true tree (*t*-value = 111.592), between the true tree length and the substitution model used for simulation (Fig. 1a; *t*-value = −104.186), and between the true tree length and stemminess of the true tree (*t*-value = −86.007).

Several of the primary factors explaining error in branch-length estimates were the same as those explaining error in the tree topology, including the interaction between true tree length and the substitution model used for simulation (Fig. 1; *t*-value = −191.171), and the interaction between tree imbalance and the stemminess of the true tree (*t*-value = 110.811). In addition, a strong predictor of error in branch-length estimates was the interaction between the substitution model used and the proportion of invariable sites (*t*-value = 215.571). All *p*-values were small (<0.001) and remained qualitatively identical when adjusted using false-discovery rates.

The entropy *t*-statistics were sensitive to factors that were similar, but not identical, to those that best explained phylogenetic error. The coefficients of regressions explaining topological error and the entropy statistics were similar (Fig. 2a, b). Meanwhile, coefficients of regression explaining error in tree-length estimate were not associated with those explaining the entropy statistics (Fig. 2c, d). Therefore, the saturation test presented here is primarily suited to assessing misleading estimates of tree topology rather than branch lengths.

**Figure 2.**
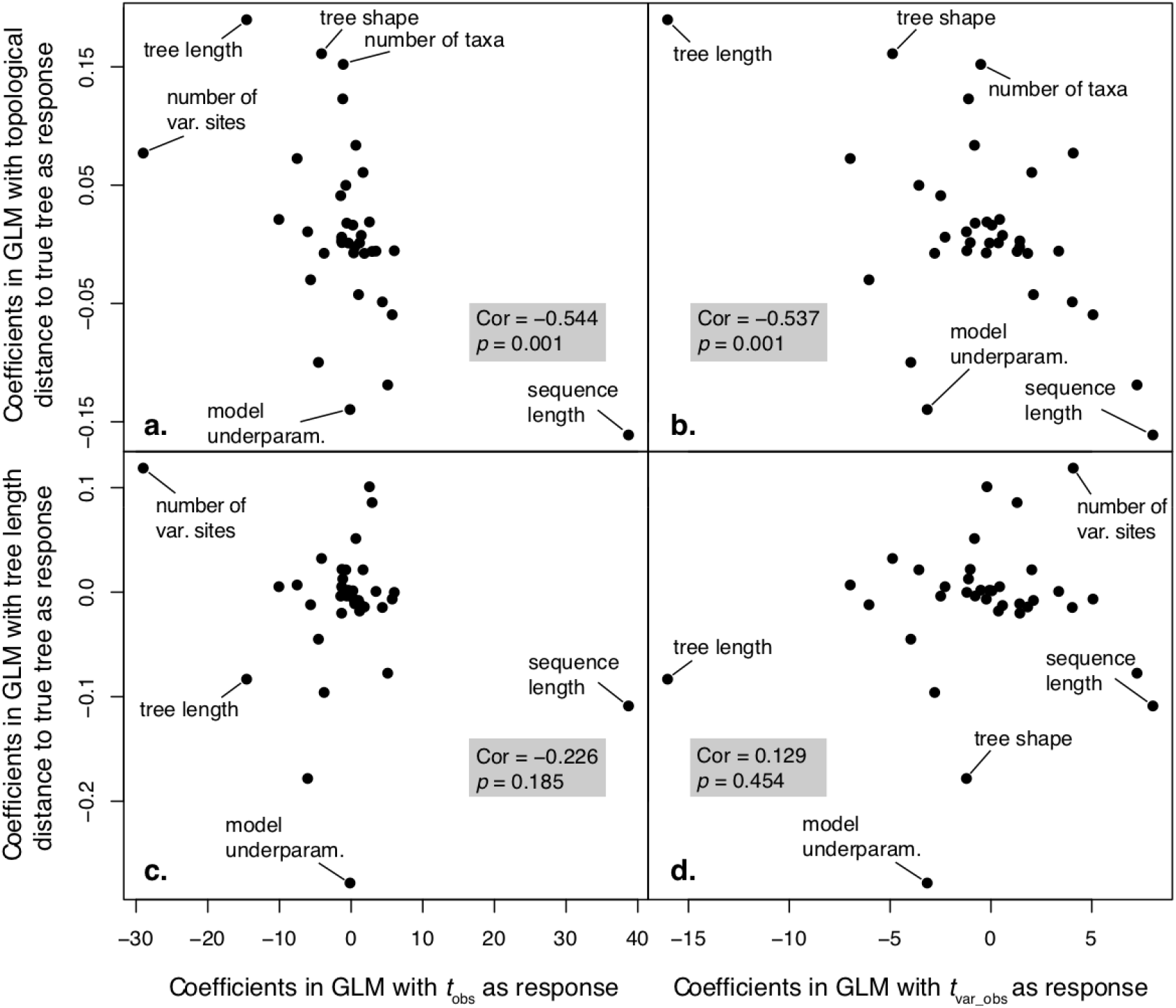
Coefficients of regression analyses of multiple variables across simulations explaining topological distance to the true tree and tree length distance (*y*-axis) and the entropy *t*-statistics (*x*-axis). A strong negative correlation between regression coefficients explaining topological distance and entropy *t*-statistics (a, b) indicates that the statistics have some power to predict misleading inferences of tree topology. Meanwhile, there is no association in regression coefficients explaining tree-length distance to the true tree and the entropy statistics (c, d).

The *t*-statistic applied to phylogenetically informative sites has a greater sensitivity to factors that also affect phylogenetic inference compared with the statistic on all sites. The statistic calculated on all sites was best explained by the interaction between true tree length and stemminess of the true tree (*t*-value = 144.484), followed by the interaction between true tree length and phylogenetic imbalance (*t*-value = 114.893). However, the test statistic is weakly correlated with an underparameterized substitution model and extreme cases of tree imbalance (Supplementary Table S1). In contrast, the statistic on phylogenetically informative sites was reasonably well explained by the interaction between the true tree length and the substitution model used for simulation (*t*-value = 124.593), and between imbalance and stemminess (*t*-value = 72.962).

### Diagnostic ability of the entropy t-statistic

The entropy *t*-statistic test focusing on informative sites consistently made a clear separation between true positives and false positives across a traverse of values of the test statistic, suggesting strong power to discriminate between accurate and misleading inferences of tree topology (Fig. 3). In contrast, the test using all alignment sites performed poorly in distinguishing true positives from false positives, and was virtually ineffective when the substitution model was underparameterized. Under a matched substitution model, the smallest number of taxa explored (8) led to the poorest test performance, primarily caused by a minority of true negatives having values that resemble most of the true positives (i.e., some accurate inferences having a signal that resembles high entropy; Fig. 3). Substitution model underparameterization primarily affected test performance in analyses with large numbers of taxa. Long sequences led to a small drop in test performance in analyses with large numbers of taxa (128 and 512 taxa).

**Figure 3.**
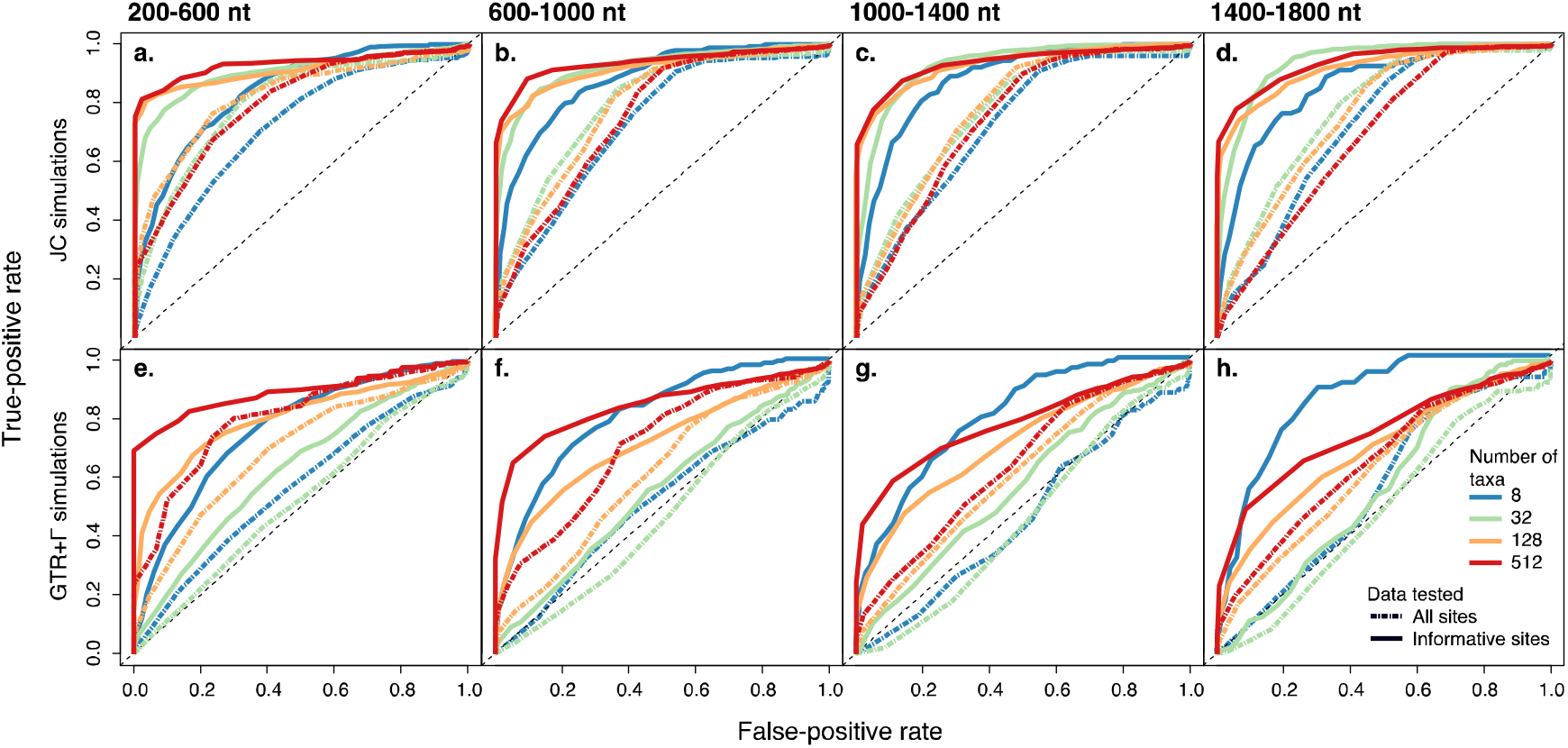
Receiver operating characteristic (ROC) curves showing the power of the entropy *t*-statistic test to classify phylogenetic topological inferences that are accurate versus inaccurate. Panel columns separate the results across four categories of alignment length, each of width 400 nt. Panel rows separate the results of analyses where the model used for simulation matched that used for inference (a–d), versus scenarios with model underparameterization (e–h). The curves show the proportion of true positives and false positives as discriminated across the range of values of the entropy *t*-statistic. Each line shows the results from a set of approximately 3,100 simulations based on random combinations of substitution rates, tree balance/imbalance, and tree stemminess.

The number of taxa and sequence length are suitable predictors of the best thresholds as taken from the data in ROC curves (the value that maximizes the difference between numbers of true positives and false positives). Specifically, according to multiple regression, the values of *t_crit_* of the test on informative sites are well explained by the square root of numbers of taxa and sequence length (adjusted *r^2^* = 0.9; Fig. 4). Nonetheless, there will be excessive uncertainty around the predicted threshold values when the number of taxa is ≫512 and sequence length is ≫1600. Another observation is the high values of *t_crit_* at intermediate numbers of taxa. At these tree sizes, estimates of base composition will be the most influenced by sites that are slow-evolving yet are parsimony-informative. For this reason, base composition is the least representative of maximum entropy at intermediate numbers of sites. This is reflected in the high variance in estimates of *t_crit_* and higher predicted values at those intermediate numbers of taxa (Fig. 4).

**Figure 4.**
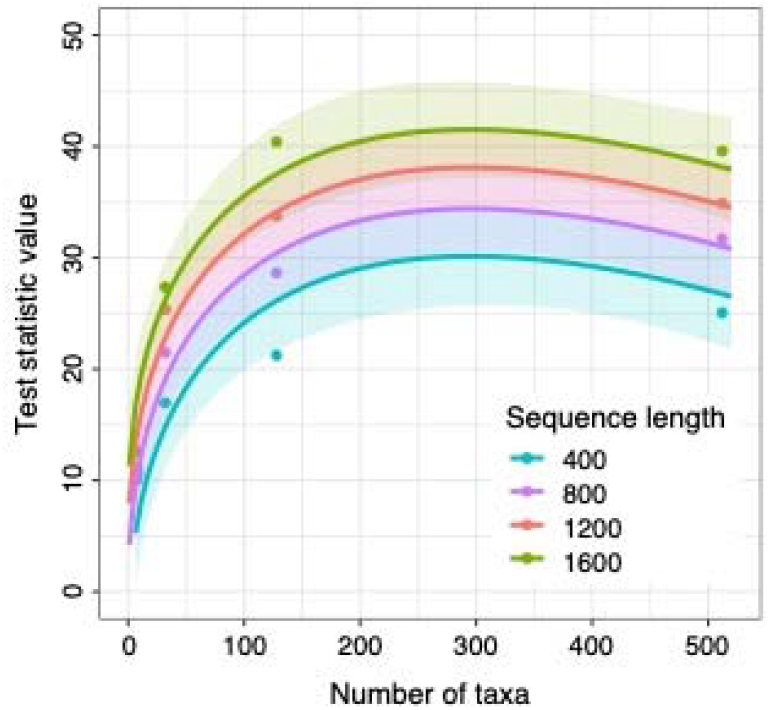
Predicted values of *t_crit_* using a multiple regression in which the explanatory variables are the square root of sequence length and number of taxa. The predictions are made for a test using exclusively phylogenetically informative sites. Each critical value is chosen to maximize the difference between numbers of true positives and false positives (taken from the data shown in Fig. 3). Lines show the predicted values of the *t_crit_* across different numbers of taxa, separated in color by each of the values of sequence length considered. Shading indicates the uncertainty around the predicted values of *t_crit_* across numbers of taxa and sequence lengths.

The test on informative sites also had a reasonable ability to identify inaccurate inferences of branch lengths (Supplementary Fig. S1), while the test on all sites had poor performance throughout scenarios. The test on informative sites was also virtually unaffected by model underparameterization when including small numbers of taxa. These results are consistent with our simulations showing that the primary drivers of misleading branch-length estimates are large tree lengths, high stemminess and excessive tree imbalance, rather than the accuracy of the substitution model among those examined.

### Signal of substitution saturation in phylogenomic data

Our implementation of the entropy tests of saturation on empirical data sets took an average of 0.141 seconds per locus. Saturation was flagged as being highly prevalent in more than a quarter of the data sets examined (10 of 36), affecting >5% of loci in these data sets and >50% in one case. In the remaining data sets, saturation was rarely flagged, or not at all (26 of 36). Given this striking distinction among data sets, our analyses show that, when prevalent, saturation can have a large impact on phylogenomic analyses. Branch support (aLRT) was on average lower in saturated than unsaturated loci in 70% of the data sets that had any flagged saturation (Fig. 5a). Unsaturated loci also had a number of informative sites that was high relative to other loci in their corresponding study (Fig. 5b). This means that rather than having an absolute small number of informative sites, it was loci with relatively small numbers of informative sites that tended to be flagged for saturation. Saturated and unsaturated loci led to similar estimates of branch lengths and stemminess, and had comparable GC contents (Supplementary Fig. S2).

**Figure 5.**
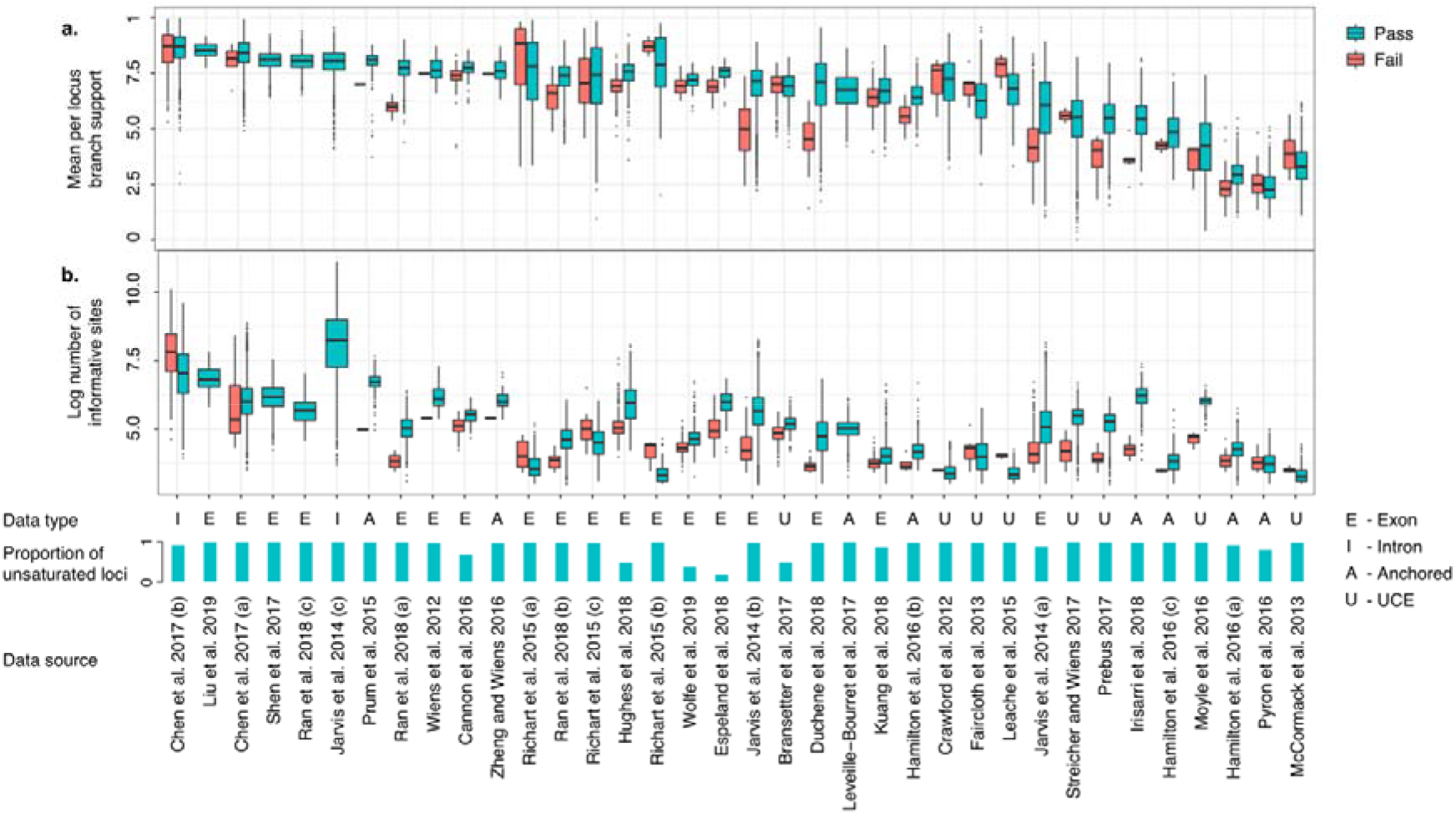
Characteristics of loci with and without substitution saturation from 36 phylogenomic data sets: (a) mean branch support in inferred gene trees, and (b) the number of informative sites across loci. Data sets are ordered from left to right by the mean branch support in trees inferred from unsaturated loci. The x-axis shows for each study the data type, the proportion of loci found to have no substitution saturation, and the source publication.

Our analyses revealed that the impact of saturation on discordance across sites in gene trees can be substantial. Loci flagged for saturation yielded gene trees that had mean aLRT branch supports that were 4.8% lower on average than the trees inferred from unsaturated loci, being 25% poorer in one data set (Fig. 6). Nonetheless, the portion of saturated loci was not associated with the loss in branch support in gene trees from saturated loci (Fig. 6b). The data sets that benefited the least from distinguishing between saturated and unsaturated loci tended to be UCE or anchored-enriched data. These data sets also had a slight tendency to have fewer informative sites per locus and smaller numbers of taxa (Fig. 6c, d).

**Figure 6.**
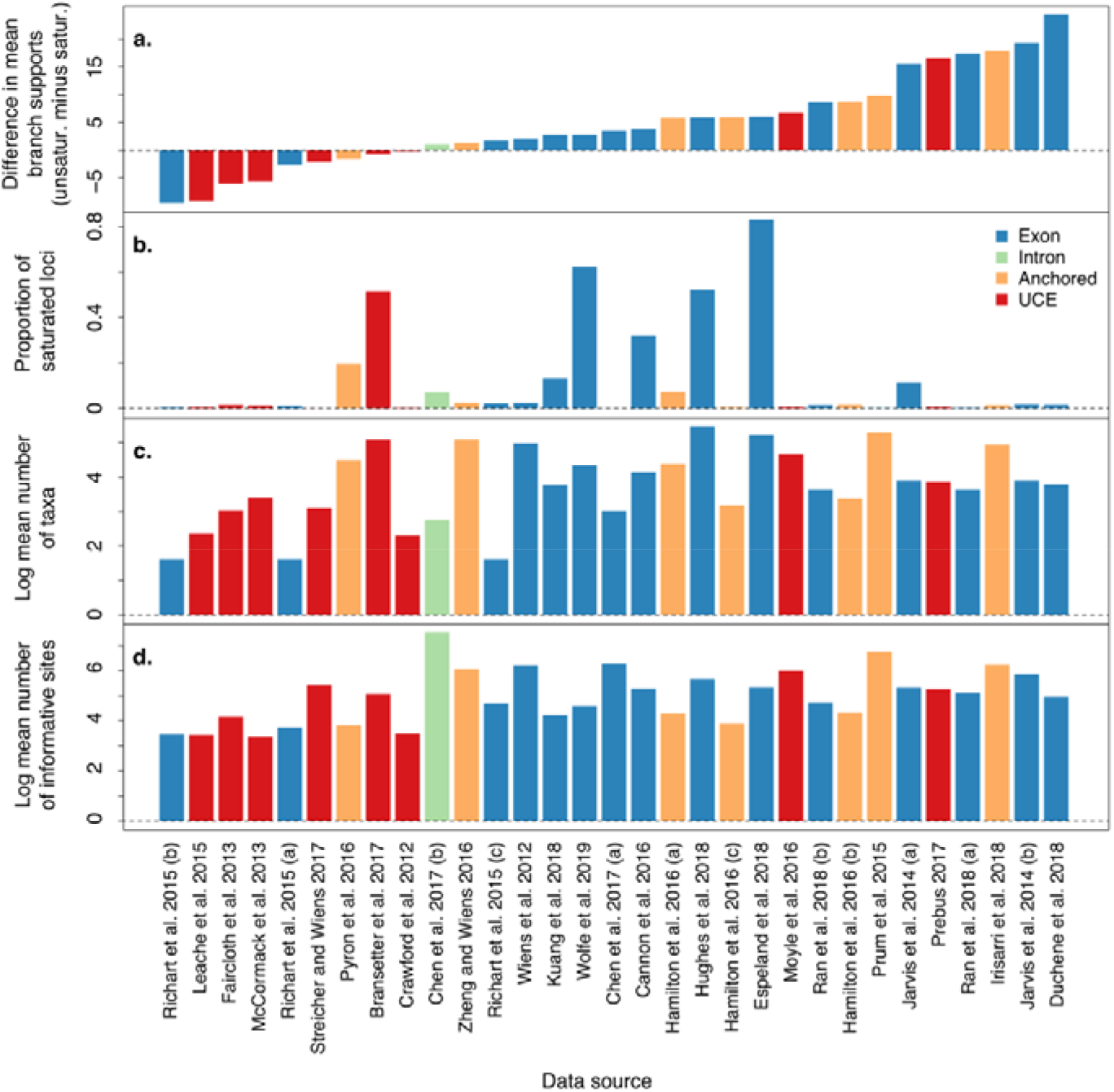
Characteristics of empirical phylogenomic data sets ranked by the difference in mean branch support between gene trees from unsaturated versus saturated loci (a). Mean branch supports are taken from mean aLRT supports (Guindon et al. 2010) across branches. The portion of saturated loci (b) is 1 minus the portion of unsaturated loci. Values of number of taxa and number of informative sites are the mean across loci in each data set.

Data sets that targeted exons and introns had higher proportions of saturated loci than those that targeted UCEs and anchored-enriched regions (Fig. 7a). Data sets from exons/introns also tended to have greater numbers of informative sites, yield longer gene trees with greater aLRT branch supports than data sets from UCEs and anchored-enriched regions (Fig. 7b, c). In addition, data sets comprising exon/introns led to gene trees with shorter internal relative to external branch lengths (lower stemminess; Fig. 7e). The longer internal branches found in gene trees inferred from UCEs or anchored-enriched regions suggest that low branch supports are likely associated with the lower number of sites and overall shorter trees, rather than due to misleading inferences such as those arising from substitution saturation (Fig. 5).

**Figure 7.**
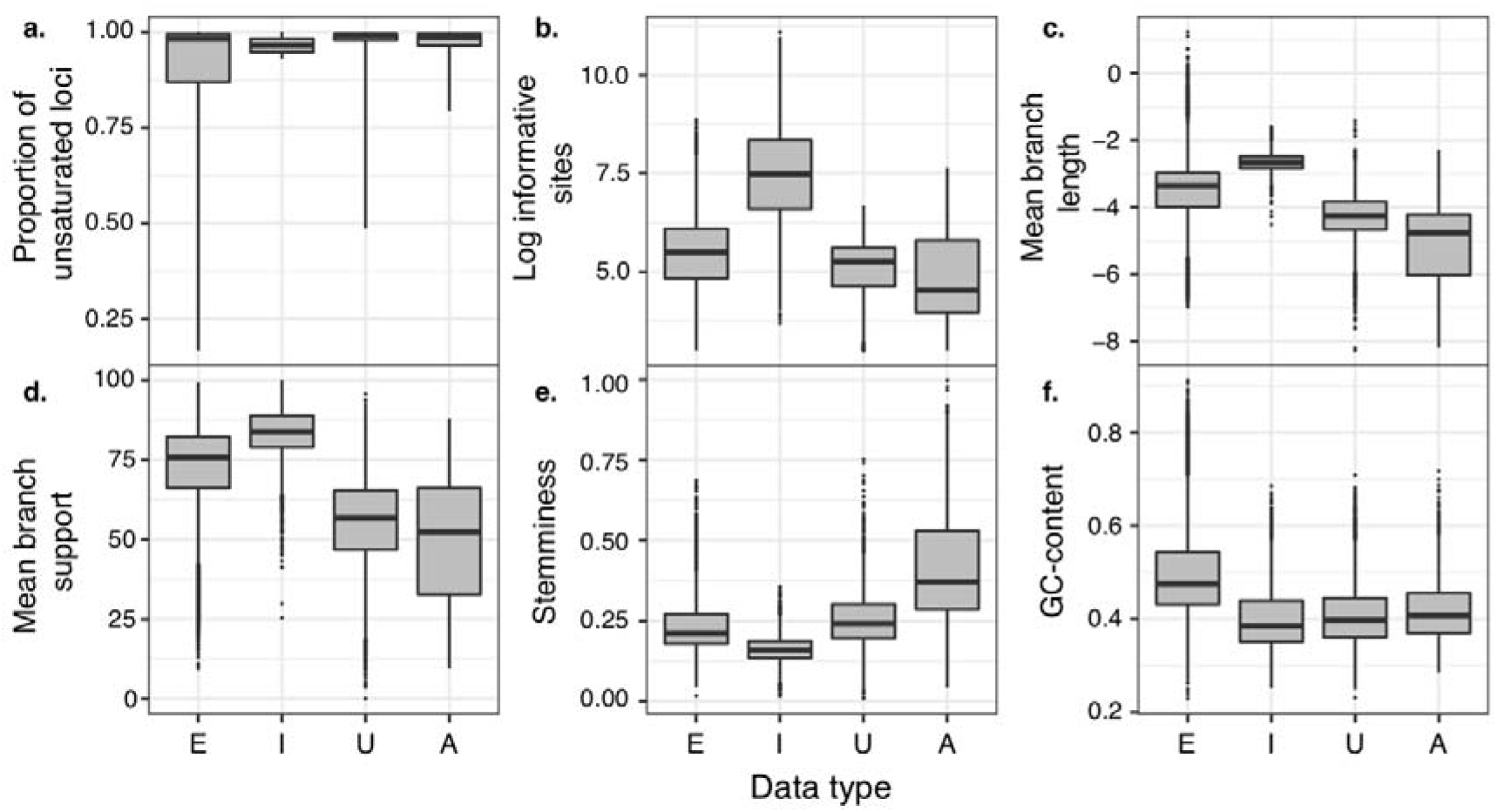
Characteristics of each data type grouped across the studies examined. Data types include exon (E), intron (I), ultraconserved elements (U), and anchored-enriched regions (A). Characteristics of data types include (a) the results from the new test of saturation as the portion of loci across studies identified as unsaturated. Data in panel (a) represent values per data set, while those in panels (b-f) show data per locus. The log number of informative sites (b) refers to sites that are parsimony-informative. Mean branch supports (c) were calculated using aLRT (Guindon et al. 2010). Stemminess (e) refers to the ratio of summed internal to external branch lengths (Fiala and Sokal 1985).

## Discussion

Our study has shown that substitution saturation can be highly misleading in phylogenetics, even when compared with a range of other factors that can also mislead inference. Saturation was flagged as occurring in large numbers of loci in nearly one-third of a broad range of empirical data sets examined, and tended to be flagged in loci with highly discordant signals across sites (Supplementary Fig. S3). Interestingly, loci with relatively few informative sites in each data set were frequently flagged as being saturated, such that they should be scrutinized in empirical data. One solution is to use the entropy *t*-statistic calculated on phylogenetically informative sites to rapidly identify levels of substitution saturation that are likely to mislead phylogenetic inference. Similar tests for identifying historical signal in sequence alignments vary considerably in their effectiveness (e.g., Strimmer and Von Haeseler 1997; Goldman 1998; Townsend 2007; Susko and Roger 2012; Townsend et al. 2012; Klopfstein et al. 2017). For this reason, we have focused on a description of the rates of true positives versus false positives of entropy tests for substitution saturation and a fast implementation. Our results indicate that saturation is a relatively common and problematic phenomenon in phylogenomics; accordingly, the entropy test of saturation as described here offers a useful complement to existing diagnostics of data quality and model performance.

Our analyses of empirical data sets suggest that different data types have comparable portions of saturated loci. Rather than depending on data type, saturation is typically flagged as a problem for loci that have small numbers of fast-evolving sites. This finding lends support to the hypothesis that variable sites in highly conserved genomic regions have more saturation than those in highly variable genome regions (Philippe et al. 1996). Despite the similar results across data types, we also observed only limited improvement in gene-tree branch supports when excluding saturated loci in UCE data sets (Supplementary Fig. S3). This might be due to a more careful choice of markers and data filtering before analyses of UCEs, which is also reflected in the small portions of loci rejected.

Strikingly, our test identified large proportions of loci as being saturated in multiple data sets, in some cases identifying saturation in more than half of the data. This suggests that assessing substitution saturation can have a dramatic effect in reducing noise in phylogenomic data sets. Assessing saturation in alignments as a whole can be particularly useful after filtering by taxa (e.g., Aberer et al. 2013; Mai and Mirarab 2018) and by sites (e.g., Ranwez et al. 2011; Whelan et al. 2018). Using a combination of methods for filtering by rate might also reduce the portion of the false negatives that arise in tests of historical signal due to fast-evolving segments of alignments (Dornburg et al. 2019).

Our findings on the performance of the entropy saturation test also point to the importance of assessing substitution model adequacy in phylogenetic studies (e.g., Goldman 1993; Bollback 2002; Weiss and von Haeseler 2003; Foster 2004; Brown 2014; Duchêne et al. 2018c). Our results from the saturation tests applied to all sites of an alignment suggest that complex evolutionary models can mislead any entropy-based saturation test on an alignment as a whole. Examples of more complex models that can mislead the test include a process in which substitution is dominated by a small number of distinct categories of base frequencies, also known for its implementation as the CAT model (Fitch and Markowitz 1970; Lartillot and Philippe 2004). Another difficult scenario is that of the covarion substitution process, where substitution types are constrained at various points in evolutionary time (Miyamoto and Fitch 1996). These models do not lead to alignment-wide base frequencies under maximum entropy, such that our saturation test is an inadequate representation of the model that generated the data. Therefore, we encourage the testing of a broad range of evolutionary models in phylogenomics. Alternatively, researchers can use multiple methods of model assessment, beginning with tests of signals in individual taxa or sites, followed by tests of model adequacy, and finishing with overall tests of historical signal (e.g., Dávalos and Perkins 2008; Liu et al. 2014).

The entropy test of substitution saturation on phylogenetically informative sites shows good performance across a broad range of scenarios, and is likely to be robust to other factors that were not explored in this study. For example, the test is likely to perform well in the presence of rate variation across sites, provided that this form of variation is modeled appropriately (Kalyaanamoorthy et al. 2017). However, due to the unusual nature of the tree-topology parameter, tests of data quality are generally dissociated from the actual performance of phylogenetic methods (also see Duchêne et al. 2017, 2018c). Outlier fast-evolving lineages might pose a challenge to tests of historical signal like the entropy saturation test, due to the possible mixed signals of closely related and highly divergent taxa (Dornburg et al. 2019). Therefore, it is useful to complement tests of the historical signal with tests of the plausibility of branch-length estimates, or of the consistency in phylogenetic signal across an alignment (e.g., Minin et al. 2003; Aberer et al. 2013; Mai and Mirarab 2018).

An important matter when developing tests of phylogenetic signal or model adequacy is to identify appropriate critical values that balance the rates of true and false positives. Identifying such critical values can be a difficult task, for many reasons. In particular, phylogenetic information is not straightforward to capture in test statistics that summarize the features of sequence alignments (Duchêne et al. 2018c). We find that the entropy saturation test is associated with several factors that affect the quality of phylogenetic inference, such as stemminess and evolutionary rate. However, the test might vary in usefulness across data sets and among loci within a phylogenomic data set. This is in part because of variance across loci in the performance of the substitution model, which will affect the performance of the test of saturation. Users of tests of model adequacy and saturation need to be aware of this limitation of the tests, and we recommend at least reporting the predicted rates of true and false positives across various data sizes from our simulations. Further work on methods of reporting the uncertainty around critical values of assessment will be valuable.

Alternative methods of assessing substitution saturation might prove to have better performance. For example, a common practice for model assessment in evolutionary biology is to use null distributions based on simulations (Brown and Thomson 2018), which allows for a test that is highly tailored to the data. However, simulations can be computationally demanding, and a simulations-based test would be dependent on using an adequate substitution model. Yet another alternative approach is to train a machine-learning algorithm to assess historical signal. Machine learning has recently been proposed in phylogenetics for substitution model selection (Abadi et al. 2020), inference of tree topology (Suvorov et al. 2019), species delimitation (Derkarabetian et al. 2019), and analyses of molecular rates across lineages (Tao et al. 2019). A random forest or an artificial neural network might prove to be highly effective for identifying the factors that are associated with accurate inferences. Similarly, these algorithms could be trained for identifying sequence alignments with misleading signals. These alternative methods might have superior performance to entropy-based tests of saturation. Nonetheless, the computational demand of the test presented here is minimal, and is unlikely to be reduced substantially using other frameworks.

Substitution saturation is detrimental to phylogenetic inference and is common in phylogenomic data sets, but it can be effectively identified using appropriate tests. Phylogenomic data sets are now widespread and researchers need to identify the data, models, and methods that are most suitable for answering the biological questions being posed (e.g., Reddy et al. 2017; Molloy and Warnow 2018; Richards et al. 2018; Bravo et al. 2019; Karin et al. 2019). The entropy test performs well across a wide range of simulation scenarios, and we provide guidelines for its usage. Tests of substitution saturation and model adequacy will improve the quality of phylogenetic inference in the genomic era, particularly in studies using data from exons or introns.

## Acknowledgements

This work was supported by funding from the Carlsbergfondet of Denmark to D.A.D. (grant CF18-0223) and S.Y.W.H (grants DP160104173 and FT160100167). The authors acknowledge the Sydney Informatics Hub and the University of Sydney’s high-performance computing cluster Artemis for providing computing resources that contributed to the research results reported in this paper.

## Notes

### Competing Interest Statement

The authors have declared no competing interest.

