## Supplementary Table S1 for "Excluding loci with substitution saturation improves inferences from phylogenomic data"

Running head : Saturation in phylogenomics

**Table S1.** Results of four linear regressions where the explanatory variables were each of the six simulation schemes, the total number of variable sites in the data, and all of the two-way interactions among them. The response variables include (a) the unweighted Robinson-Foulds distance between the estimated and true tree topologies, (b) the difference in total tree length between estimated and true tree topologies, calculated as (estimated - true) / true summed branch lengths, (c) the entropy *t*-statistic calculated from all alignment sites in each simulated alignment, and (d) the entropy *t*-statistic calculated from the phylogenetically informative sites in each alignment. Regression parameters with *p*-values < 0.001 are shown in bold.

| **a. Unweighted Robinson-Foulds distance from estimated to true tree topology** | | | |
| --- | --- | --- | --- |
|  | Estimate | Standard error | *t-*value |
| **(Intercept)** | **0.394** | **0.002** | **242.464** |
| **Substitution model** | **-0.14** | **0.001** | **-125.38** |
| **Tree length** | **0.19** | **0.001** | **224.952** |
| **Number of taxa** | **0.152** | **0.001** | **177.821** |
| **Sequence length** | **-0.161** | **0.008** | **-19.846** |
| **Tree imbalance** | **0.161** | **0.001** | **208.573** |
| **Proportion of invariable sites** | **0.011** | **0.001** | **7.282** |
| **Stemminess** | **-0.03** | **0.001** | **-38.814** |
| **Total number of variable sites** | **0.077** | **0.008** | **9.252** |
| **Substitution model : Tree length** | **-0.119** | **0.001** | **-104.186** |
| **Substitution model : Number of taxa** | **0.006** | **0.001** | **5.34** |
| Substitution model : Sequence length | -0.006 | 0.005 | -1.229 |
| Substitution model : Tree imbalance | -0.002 | 0.001 | -2.086 |
| **Substitution model : Proportion of invariable sites** | **0.019** | **0.001** | **14.335** |
| **Substitution model : Stemminess** | **-0.042** | **0.001** | **-38.763** |
| Substitution model : Total number of variable sites | -0.008 | 0.005 | -1.534 |
| **Tree length : Number of taxa** | **-0.008** | **0.001** | **-13.368** |
| **Tree length : Sequence length** | **-0.1** | **0.004** | **-24.991** |
| **Tree length : Tree imbalance** | **-0.006** | **0.001** | **-10.15** |
| **Tree length : Proportion of invariable sites** | **0.008** | **0.001** | **9.321** |
| **Tree length : Stemminess** | **-0.049** | **0.001** | **-86.007** |
| **Tree length : Total number of variable sites** | **0.084** | **0.004** | **21.078** |
| Number of taxa : Sequence length | 0.001 | 0.005 | 0.233 |
| **Number of taxa : Tree imbalance** | **0.123** | **0.001** | **218.971** |
| Number of taxa : Proportion of invariable sites | 0.001 | 0.001 | 1.174 |
| **Number of taxa : Stemminess** | **0.016** | **0.001** | **28.871** |
| Number of taxa : Total number of variable sites | 0.001 | 0.005 | 0.293 |
| **Sequence length : Tree imbalance** | **-0.059** | **0.002** | **-25.077** |
| Sequence length : Proportion of invariable sites | -0.005 | 0.003 | -2.018 |
| Sequence length : Stemminess | 0.003 | 0.002 | 1.214 |
| **Sequence length : Total number of variable sites** | **0.041** | **0.001** | **29.305** |
| **Tree imbalance : Proportion of invariable sites** | **0.018** | **0.001** | **27.511** |
| **Tree imbalance : Stemminess** | **0.061** | **0.001** | **111.592** |
| **Tree imbalance : Total number of variable sites** | **0.073** | **0.002** | **30.099** |
| **Proportion of invariable sites : Stemminess** | **-0.007** | **0.001** | **-11.181** |
| **Proportion of invariable sites : Total number of variable sites** | **0.021** | **0.003** | **7.178** |
| **Stemminess : Total number of variable sites** | **0.05** | **0.002** | **20.72** |
| **b. Summed branch length difference between estimated and true trees** | | | |
| **(Intercept)** | **-0.171** | **0.001** | **-296.586** |
| **Substitution model** | **-0.278** | **0** | **-703.114** |
| **Tree length** | **-0.083** | **0** | **-277.524** |
| **Number of taxa** | **0.002** | **0** | **6.287** |
| **Sequence length** | **-0.109** | **0.003** | **-37.73** |
| **Tree imbalance** | **0.032** | **0** | **117.444** |
| **Proportion of invariable sites** | **-0.178** | **0.001** | **-344.2** |
| **Stemminess** | **-0.012** | **0** | **-43.849** |
| **Total number of variable sites** | **0.119** | **0.003** | **40.075** |
| **Substitution model : Tree length** | **-0.078** | **0** | **-191.171** |
| **Substitution model : Number of taxa** | **0.005** | **0** | **12.987** |
| **Substitution model : Sequence length** | **0.086** | **0.002** | **49.326** |
| **Substitution model : Tree imbalance** | **-0.011** | **0** | **-28.475** |
| **Substitution model : Proportion of invariable sites** | **0.101** | **0** | **215.571** |
| **Substitution model : Stemminess** | **-0.008** | **0** | **-20.321** |
| **Substitution model : Total number of variable sites** | **-0.096** | **0.002** | **-54.618** |
| **Tree length : Number of taxa** | **-0.014** | **0** | **-69.336** |
| **Tree length : Sequence length** | **-0.045** | **0.001** | **-31.697** |
| **Tree length : Tree imbalance** | **0.001** | **0** | **3.544** |
| **Tree length : Proportion of invariable sites** | **-0.013** | **0** | **-44.159** |
| **Tree length : Stemminess** | **-0.015** | **0** | **-72.682** |
| **Tree length : Total number of variable sites** | **0.051** | **0.001** | **36.352** |
| **Number of taxa : Sequence length** | **-0.018** | **0.002** | **-10.117** |
| **Number of taxa : Tree imbalance** | **0.013** | **0** | **63.432** |
| **Number of taxa : Proportion of invariable sites** | **0.002** | **0** | **5.58** |
| **Number of taxa : Stemminess** | **0.001** | **0** | **6.767** |
| **Number of taxa : Total number of variable sites** | **0.022** | **0.002** | **12.401** |
| **Sequence length : Tree imbalance** | **-0.007** | **0.001** | **-7.799** |
| Sequence length : Proportion of invariable sites | 0 | 0.001 | -0.178 |
| **Sequence length : Stemminess** | **-0.02** | **0.001** | **-23.871** |
| **Sequence length : Total number of variable sites** | **-0.004** | **0** | **-7.513** |
| **Tree imbalance : Proportion of invariable sites** | **-0.004** | **0** | **-15.852** |
| **Tree imbalance : Stemminess** | **0.021** | **0** | **110.811** |
| **Tree imbalance : Total number of variable sites** | **0.007** | **0.001** | **8.059** |
| **Proportion of invariable sites : Stemminess** | **-0.007** | **0** | **-28.816** |
| **Proportion of invariable sites : Total number of variable sites** | **0.005** | **0.001** | **5.054** |
| **Stemminess : Total number of variable sites** | **0.021** | **0.001** | **25.029** |
| **c. Entropy *t*-statistic calculated on all alignment sites** | | | |
| **(Intercept)** | **28.443** | **0.087** | **328.685** |
| Substitution model | -0.156 | 0.059 | -2.622 |
| **Tree length** | **-14.584** | **0.045** | **-324.349** |
| **Number of taxa** | **-1.103** | **0.046** | **-24.205** |
| **Sequence length** | **38.708** | **0.433** | **89.474** |
| **Tree imbalance** | **-4.126** | **0.041** | **-100.248** |
| **Proportion of invariable sites** | **-6.066** | **0.078** | **-78.118** |
| **Stemminess** | **-5.657** | **0.041** | **-137.457** |
| **Total number of variable sites** | **-29.012** | **0.444** | **-65.344** |
| **Substitution model : Tree length** | **5.101** | **0.061** | **83.914** |
| **Substitution model : Number of taxa** | **-1.319** | **0.06** | **-21.832** |
| **Substitution model : Sequence length** | **2.928** | **0.261** | **11.241** |
| **Substitution model : Tree imbalance** | **0.498** | **0.058** | **8.534** |
| **Substitution model : Proportion of invariable sites** | **2.555** | **0.07** | **36.426** |
| **Substitution model : Stemminess** | **1.033** | **0.058** | **17.691** |
| **Substitution model : Total number of variable sites** | **-3.772** | **0.263** | **-14.32** |
| **Tree length : Number of taxa** | **1.849** | **0.031** | **60.547** |
| **Tree length : Sequence length** | **-4.559** | **0.213** | **-21.432** |
| **Tree length : Tree imbalance** | **3.466** | **0.03** | **114.893** |
| **Tree length : Proportion of invariable sites** | **1.412** | **0.043** | **32.679** |
| **Tree length : Stemminess** | **4.359** | **0.03** | **144.484** |
| Tree length : Total number of variable sites | 0.661 | 0.212 | 3.124 |
| **Number of taxa : Sequence length** | **1.181** | **0.268** | **4.409** |
| **Number of taxa : Tree imbalance** | **-1.166** | **0.03** | **-38.992** |
| **Number of taxa : Proportion of invariable sites** | **-0.383** | **0.049** | **-7.745** |
| **Number of taxa : Stemminess** | **0.242** | **0.03** | **8.081** |
| **Number of taxa : Total number of variable sites** | **-1.279** | **0.264** | **-4.844** |
| **Sequence length : Tree imbalance** | **5.73** | **0.126** | **45.331** |
| **Sequence length : Proportion of invariable sites** | **6.023** | **0.144** | **41.876** |
| **Sequence length : Stemminess** | **-1.325** | **0.126** | **-10.499** |
| **Sequence length : Total number of variable sites** | **-1.452** | **0.075** | **-19.417** |
| **Tree imbalance : Proportion of invariable sites** | **-0.607** | **0.035** | **-17.511** |
| **Tree imbalance : Stemminess** | **1.662** | **0.029** | **57.283** |
| **Tree imbalance : Total number of variable sites** | **-7.537** | **0.128** | **-58.731** |
| **Proportion of invariable sites : Stemminess** | **0.356** | **0.035** | **10.288** |
| **Proportion of invariable sites : Total number of variable sites** | **-10.079** | **0.156** | **-64.607** |
| **Stemminess : Total number of variable sites** | **-0.741** | **0.128** | **-5.782** |
| **d. Entropy *t*-statistic calculated on phylogenetically informative sites** | | | |
| **(Intercept)** | **29.623** | **0.083** | **357.912** |
| **Substitution model** | **-3.162** | **0.057** | **-55.708** |
| **Tree length** | **-16.038** | **0.043** | **-372.933** |
| **Number of taxa** | **-0.506** | **0.044** | **-11.599** |
| **Sequence length** | **8.031** | **0.414** | **19.408** |
| **Tree imbalance** | **-4.882** | **0.039** | **-124.019** |
| **Proportion of invariable sites** | **-1.215** | **0.074** | **-16.358** |
| **Stemminess** | **-6.049** | **0.039** | **-153.679** |
| **Total number of variable sites** | **4.08** | **0.425** | **9.608** |
| **Substitution model : Tree length** | **7.244** | **0.058** | **124.593** |
| **Substitution model : Number of taxa** | **-2.277** | **0.058** | **-39.392** |
| **Substitution model : Sequence length** | **1.297** | **0.249** | **5.205** |
| **Substitution model : Tree imbalance** | **1.424** | **0.056** | **25.487** |
| Substitution model : Proportion of invariable sites | -0.2 | 0.067 | -2.977 |
| **Substitution model : Stemminess** | **2.107** | **0.056** | **37.724** |
| **Substitution model : Total number of variable sites** | **-2.792** | **0.252** | **-11.082** |
| **Tree length : Number of taxa** | **1.822** | **0.029** | **62.372** |
| **Tree length : Sequence length** | **-3.975** | **0.203** | **-19.535** |
| **Tree length : Tree imbalance** | **3.359** | **0.029** | **116.421** |
| **Tree length : Proportion of invariable sites** | **0.574** | **0.041** | **13.892** |
| **Tree length : Stemminess** | **4.034** | **0.029** | **139.798** |
| **Tree length : Total number of variable sites** | **-0.816** | **0.202** | **-4.035** |
| Number of taxa : Sequence length | 0.372 | 0.256 | 1.452 |
| **Number of taxa : Tree imbalance** | **-1.114** | **0.029** | **-38.925** |
| Number of taxa : Proportion of invariable sites | -0.07 | 0.047 | -1.489 |
| Number of taxa : Stemminess | 0.049 | 0.029 | 1.707 |
| **Number of taxa : Total number of variable sites** | **-1.027** | **0.253** | **-4.066** |
| **Sequence length : Tree imbalance** | **5.06** | **0.121** | **41.854** |
| **Sequence length : Proportion of invariable sites** | **-1.196** | **0.138** | **-8.697** |
| **Sequence length : Stemminess** | **1.432** | **0.121** | **11.856** |
| **Sequence length : Total number of variable sites** | **-2.492** | **0.072** | **-34.848** |
| **Tree imbalance : Proportion of invariable sites** | **-0.784** | **0.033** | **-23.655** |
| **Tree imbalance : Stemminess** | **2.024** | **0.028** | **72.962** |
| **Tree imbalance : Total number of variable sites** | **-6.979** | **0.123** | **-56.853** |
| **Proportion of invariable sites : Stemminess** | **-0.231** | **0.033** | **-6.985** |
| Proportion of invariable sites : Total number of variable sites | 0.431 | 0.149 | 2.889 |
| **Stemminess : Total number of variable sites** | **-3.574** | **0.123** | **-29.145** |

**Figure S1.** Performance of the entropy saturation test for identifying misleading branch-length estimates. A positive was chosen to be a case where the summed of estimated branch lengths was 50% above or below of the summed simulated (true) branch lengths. The simulations include 10^5^ random draws from six simulation variables, such that each line shown represents approximately 3,100 simulations.

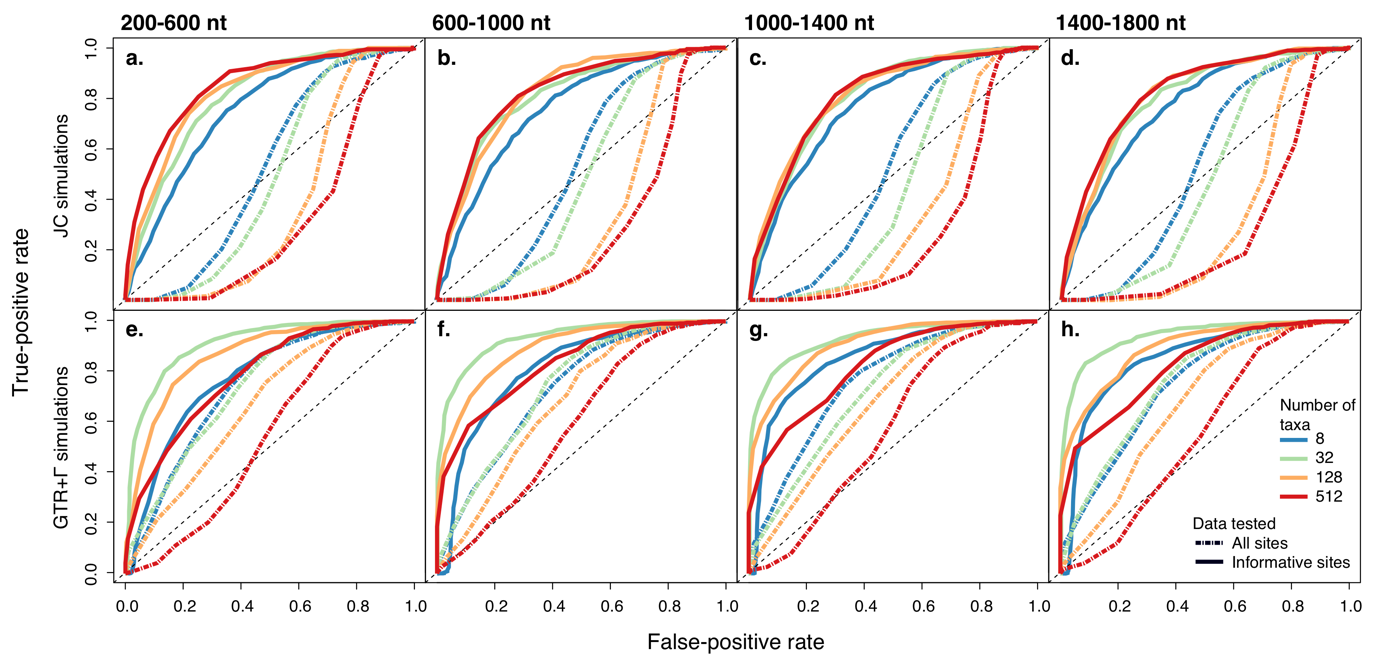

**Figure S2.** Characteristics of loci with and without saturation for each of 36 phylogenomic data sets examined. Data sets are ordered from left to right by overall branch support of non-saturated loci. The characteristics shown include (a) the log mean of branch lengths across estimated gene trees in each data set, (b) the total number of sites per locus alignment, (c) the proportion of the total tree length that is contributed by internal branches (stemminess; Fiala and Sokal 1985), and (d) the summed proportion of guanine and cytosine nucleotides across alignments in each data set. The x-axis shows the source study for each data set.

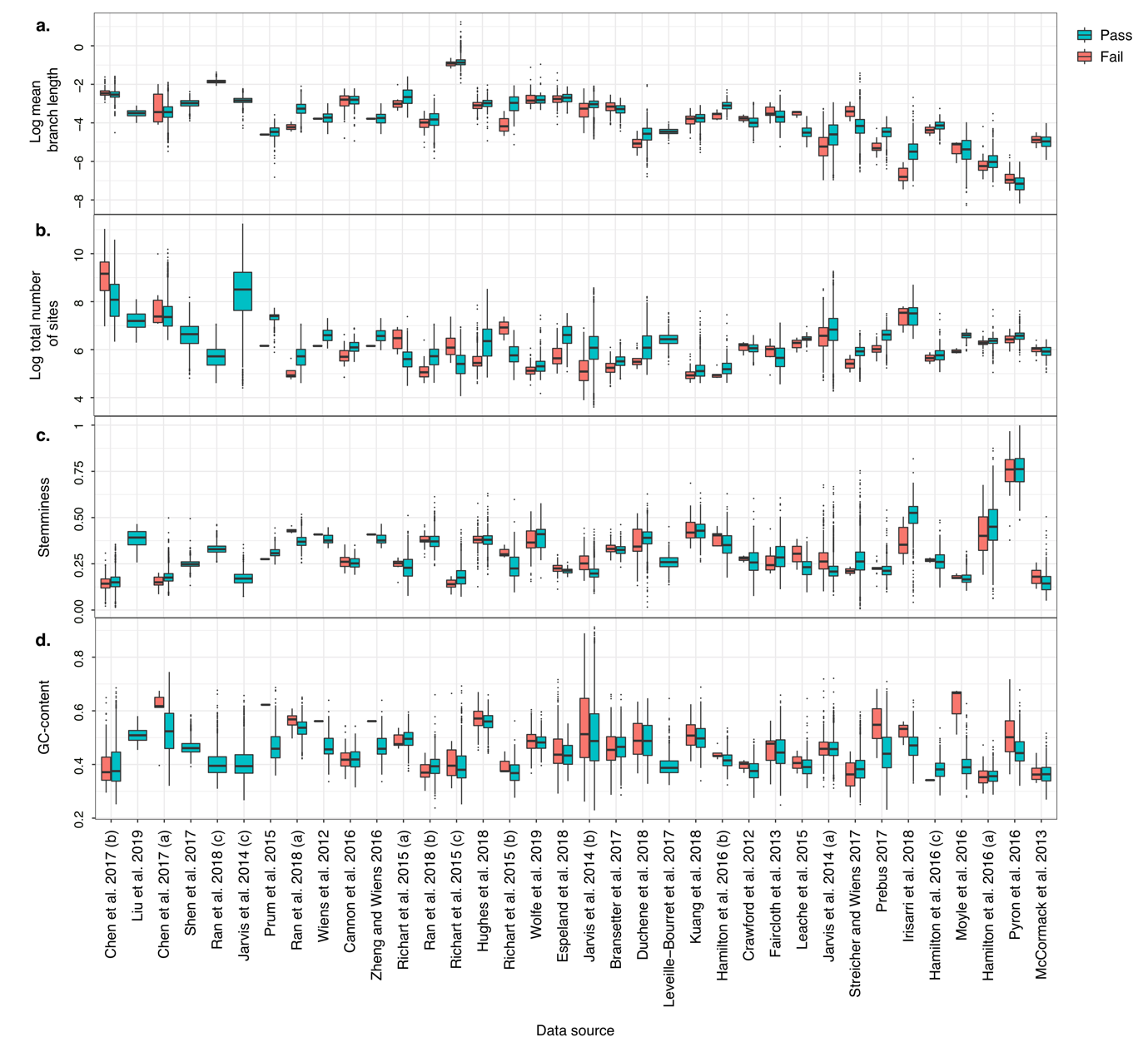

**Figure S3.** Characteristics of data sets ranked by the difference in mean branch support in estimated gene trees between unsaturated and saturated loci (see Fig. 6 in main text). Values shown are the mean across loci in each data set.

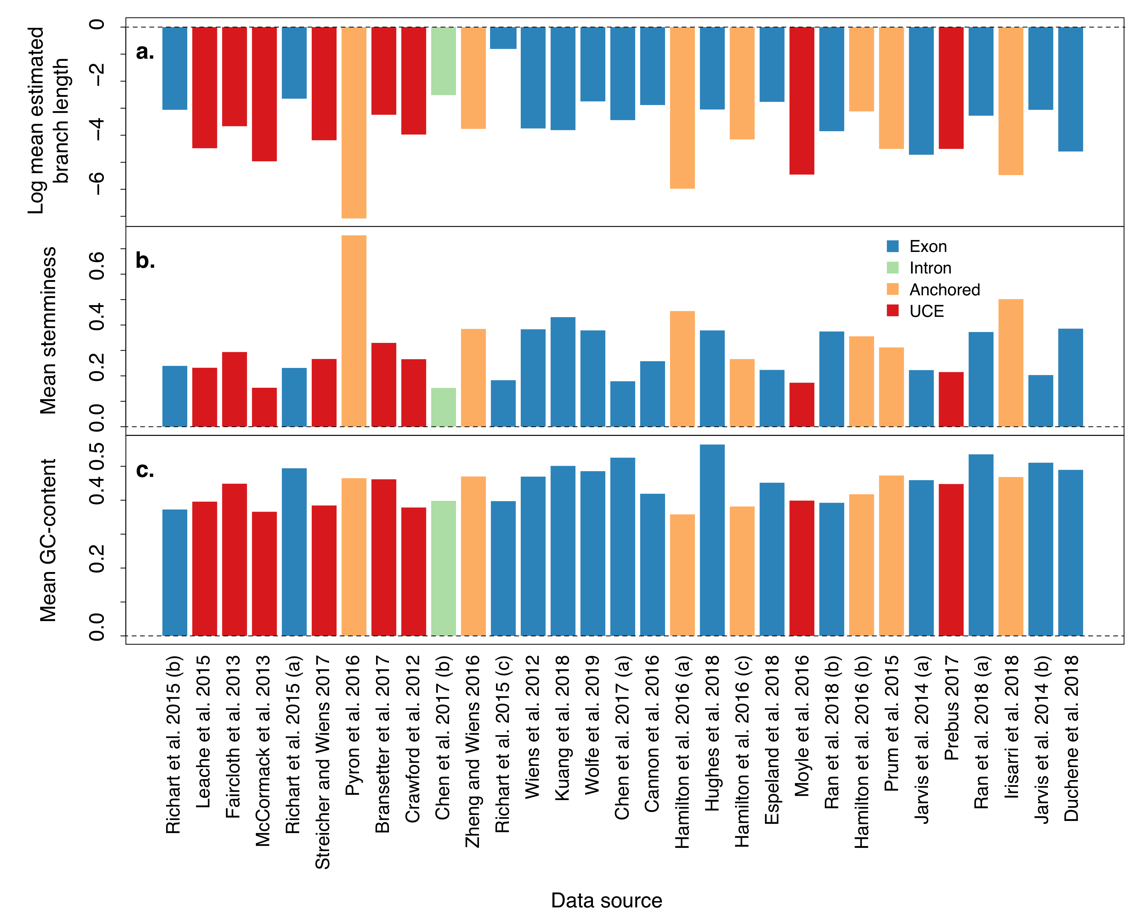
